# PELSA-Decipher: a software tool for the processing and interpretation of ligand-protein interaction dataset acquired by PELSA

**DOI:** 10.1101/2025.02.27.640683

**Authors:** Haiyang Zhu, Keyun Wang, Kejia Li, Zheng Fang, Jiahua Zhou, Mingliang Ye

## Abstract

Knowledge of protein-ligand interactions is essential for comprehending various aspects of life science, including drug mechanisms of action, regulatory processes in cellular metabolism and signaling, etc. Recently, a robust ligand modification-free method, termed as peptide-centric local stability assay (PELSA), is developed to identify the protein targets and binding regions of diverse ligands at proteomics scale. This method has unprecedented sensitivity and can be broadly applied to ligands including drugs, metabolites, metal ions, antibodies tec. However, the extraction of key information on ligand-protein interaction, including the binding protein, binding site, and binding affinity, is not a trivial task, which hampers the wide utilization of PELSA strategy by the research community. To address this, we developed a software tool, PELSA-Decipher, to facilitate the efficient processing of dataset obtained in PELSA experiment, including raw data processing, result visualization, and the generation of reports and high-quality images, thereby will greatly promote the broader applications of PELSA strategy. The PELSA-Decipher software can be downloaded free of charge from the website https://github.com/DICP-1809.

## Introduction

Proteins are the essential molecules in living organisms, and their structure is critical for their functional roles in regulating biological processes[1-3]. The functional roles of a protein always involve interactions with small molecules, including metal ions, metabolites, signaling molecules or drugs. Therefore, identifying proteins that can be modulated by specific ligands is crucial to dissect the biological function of a protein as well as the action mechanism of ligands. With the development of high-throughput screening technology, the mass spectrometry (MS) based proteomics now allows large-scale functional analysis of protein-ligand interactions[4-7].

A series of energetics-based modification-free methods have been developed by measuring the ligand induced change in specific properties of proteins. In comparison to chemical proteomics that requires the chemical modification of the ligand to synthesize probe, the energetics-based modification-free methods offers a more accurate and reliable means for ligand target identification[8-10]. For instance, ligand binding increases the stability of the target proteins, and makes them more resistant to thermal-induced protein denaturation, which enabled the development of techniques such as thermal proteome profiling (TPP)[11], isothermal shift assay (iTSA)[12], and matrix thermal shift assay (mTSA) for target protein identification[13]. Similarly, the ligand stabilized proteins are resistant to solvent-induced protein precipitation, and so led to the development of methods such as solvent-induced protein precipitation (SIP)[14], solvent proteome profiling (SPP)[15], and isosolvent shift assay (iSSA)[16], pH-dependent protein precipitation (pHDPP)[17] and iso-pH shift assay (ipHSA)[16]. Moreover, combining ipHSA, iTSA, and iSSA has resulted in the development of the integrated protein solubility shift analysis (IPSSA) method[16]. However, all above methods identify target proteins by detecting the overall change in the stability of proteins, and are unable to provide binding site information.

Ligand binding can induce local changes in protein conformation or thermodynamic property, resulting in alterations in proteolysis susceptibility, and can therefore be probed by limited proteolysis. Feng et al. proposed a two-step digestion strategy, limited proteolysis coupled with mass spectrometry (LiP-MS)[18], which enables binding region determination by assessing proteolysis susceptibility at the peptide level. However, a subsequent study revealed that LiP-MS could not confidently identify target proteins in the complex human cell lysates; Therefore, Piazza et al. developed LiP-Quant[19], a revised version of LiP-MS that requires additional ligand dose-response profiles and a machine-learning framework. However, the target protein identification capacity of LiP-Quant evaluated by staurosporine, a pan-kinase inhibitor, is much less than the prevalent modification-free method TPP. Recently, we proposed to utilize extensive trypsinization to directly generate MS-detectable peptides from native proteins to monitor protein local stability. This digestion scheme in coupled with a simple FASP procedure significantly reduces the complexity of peptide samples and can amplify the readout of ligand-induced protein local stability shifts. Based on this, we established a method, PEptide-centric Local Stability Assay (PELSA), that enables sensitive identification of target proteins as well as corresponding binding-region information. PELSA was demonstrated to have unprecedented sensitivity in identifying ligand-binding proteins and can be applied for sensitive probing of weak interactions, as exemplified by identifying the binding proteins of leucine, folate, αKG, and R2HG. Take αKG as an example, PELSA results consistently exhibit a significantly higher percentage of known-binding events. For instance, in a prior LiP-MS study of αKG-treated E.coli lysates, 34 candidate targets were identified with 2 known αKG binding proteins. In comparison, PELSA identified 40 candidate targets, and notably, 30 of them were known αKG binding proteins, despite using a more complex lysate sample (human HeLa cell lysate). [20]

In the modification-free experiments, peptides from complex cell samples are analyzed by LC-MS/MS, which will generate huge volume of raw dataset. The extraction of the ligand target interaction information from such datasets can be time-consuming and requires considerable expertise, which poses great challenges to the broader application of these methods. Therefore, streamlining and optimizing the downstream data processing workflows to enhance their efficiency and user-friendliness is crucial to promote the method’s widespread application. Numerous software tools have been developed to accommodate corresponding experimental methods, such as ProSAP[21], CHalf[22], IMPRINTS.CETSA.app[23], FLiPPR[24] and Lip-Quant[19]. These software tools and methods are capable of processing the corresponding experimental data. However, all of them are not applicable to the processing of PELSA data.

To improve the efficiency and usability of the PELSA method for target identification, we developed PELSA-Decipher, a tool designed to streamline the end-to-end processing of PELSA data. PELSA-Decipher is an efficient and user-friendly software that integrates functions such as differential analysis, protein local stability analysis, and concentration-dependent local affinity analysis **(Figure 1)**. By incorporating domain information from the UniProt database[25], the software enables precise analysis of ligand-protein binding regions and also supports user-defined regions for targeted region analysis. PELSA-Decipher offers flexible image customization features, supports multiple output formats, and allows for batch exporting of result reports and images. PELSA-Decipher is free for academic using and can be downloaded from GitHub: https://github.com/DICP-1809.

**Figure 1.**
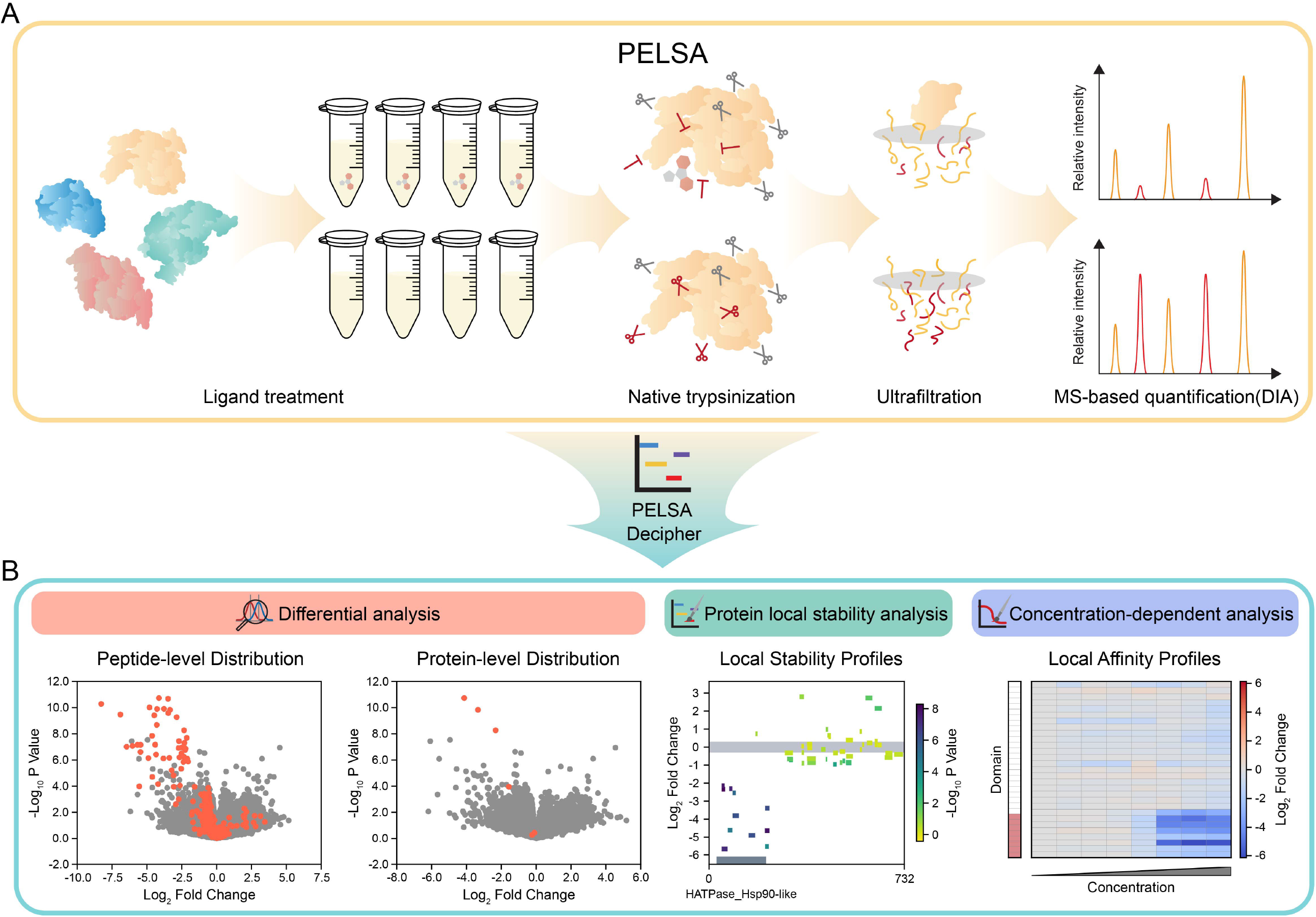
Schematics of the PELSA-Decipher. A. Workflow of PELSA. B. The three main modules of PELSA-Decipher: differential analysis to identify target proteins, protein local stability analysis to determine ligand binding regions, and concentration-dependent analysis to evaluate local and global binding affinity.

## Method

### Implementation of PELSA-Decipher

PELSA-Decipher is implemented in Python 3.11 (v3.11.9) and R (v4.3.2). By integrating the strengths of Python and R—Python’s efficiency in data manipulation and the extensive suite of robust R packages—PELSA-Decipher ensures both efficiency and reliability in the analytical workflow. Python primarily facilitates software framework construction, data preprocessing, and results visualization. The graphical user interface (GUI) is developed primarily with PySide6 (v6.7.2) and further enhanced using PySide6-Fluent-Widgets (v1.5.7). Data filtering and preprocessing are performed using Numpy (v2.0.0) and Pandas (v2.2.2), while various visualizations such as volcano profiles, local stability profiles, and local affinity profiles, are generated using Matplotlib (v3.9.2) and Seaborn (v0.13.2). R is primarily employed for data processing, requiring the preinstallation of R and its associated packages, which can be facilitated using an included installation script. The R package limma (v3.58.1)[26] is utilized for differential data analysis, and drc (v3.0-1) is applied to model concentration-dependent curves. Data exchange between Python and R is facilitated via the Rpy2 package (v3.5.16). PELSA-Decipher primarily accepts input data in CSV format. Plain text files, generated and exported by protein quantification software such as Spectronaut, DIA-NN, can also be used as the input format for PELSA-Decipher analysis. Protein domain information utilized by PELSA-Decipher is primarily downloaded from the UniProt database as TSV-format files and imported as the domain database. Additionally, JSON files containing protein domain information are supported. Users have the flexibility to customize domain information by using the built-in domain management tool to add, delete, modify, or query the database, with the option to export the data in JSON format.

### Dissecting differences in protein changes

PELSA-Decipher comprises three main modules: differential analysis (DA), protein local stability analysis (ProLSA), and concentration-dependent analysis (CDA), encompassing the entire workflow from data preprocessing and processing to result visualization and results outputting.

*In the differential analysis module*, a standardized data processing pipeline is provided, which includes data cleaning, missing value handling, coefficient of variation (CV) calculation, data transformation, and subsequent statistical analysis. Data cleaning ensures the removal of contaminant proteins and guarantees the unique mapping of peptide sequences to proteins, with the process being based on the specified identifiers. Several options are available for missing value handling, such as deletion, mean imputation, and linear interpolation, with deletion being the default. Data transformation converts the data to a different value scale, with the default being base-2 logarithm, making the data more centralized and approximating a normal distribution, thereby facilitating downstream analysis. For statistical analysis, the robust and widely used empirical Bayes method from the limma package is employed to ensure reliable and stable results. The differential expression results for proteins are represented by the peptide with the smallest p value among all peptides associated with the protein.

*The protein local stability analysis module* is designed to assess and visualize information associated with changes in protein local stability after the ligand binding. The protein domain data is obtained from the UniProt database and imported in either TSV or JSON format. The protein local stability profile utilizes the protein amino acid sequence as the x-axis, log_2_ fold change as the y-axis, and -log_10_ p-values as the color gradient, effectively illustrating the extent of protein variation across different domains. In screening approaches based on enzyme cleavage, the binding of ligands to proteins impedes the cleavage process, resulting in a decrease in the intensity of corresponding peptides during mass spectrometry quantification. Therefore, the local stability profiles effectively highlight the binding regions of ligands on the protein.

*The concentration-dependent analysis module* calculates the EC50 values at both peptide and protein levels by fitting a four-parameter equation **(Function 1)**. This module employs a standardized data processing workflow to analyze and filter results from different concentration points, outputting concentration-dependent calculation results and corresponding concentration-dependent curves. Initially, the data undergoes cleaning, interpolation, and outlier removal, followed by the calculation of CV values. Subsequently, preliminary filtering is performed based on CV values, with a default threshold set at 0.5 to exclude data exhibiting excessive variability. Next, the concentration-dependent curve at the peptide level is fitted using the average intensity of each concentration point as a percentage relative to the vehicle group. The fitted results are further scrutinized, focusing on data from the highest and second-highest concentration points, which are compared to the vehicle group for differential expression analysis. Peptides with p-values less than 0.01 (or -log_10_ p value greater than 2) are annotated as significant. The filtering criteria applied to the peptides include: 1) an R^2^ value greater than 0.9 for the concentration-response curve fitting; 2) the average intensity at the maximum concentration point being less than 0.75 relative to the vehicle group; and 3) peptides marked as significant in the differential analysis. Ultimately, the average values of the peptides that meet the filtering criteria are taken as the protein intensity, and a four-parameter equation is used for fitting to derive the concentration-dependent information for the proteins.

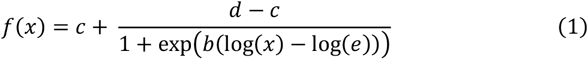

Furthermore, PELSA-Decipher enabes the definition of custom domain intervals, allowing for targeted monitoring of specific protein regions. The *Domain Assistant tool*, integrated into the PELSA-Decipher toolbox, enables users to add, delete, modify, and export domain region information to construct a customized domain database for generating protein-specific local stability profiles.

## Results

### Workflow of PELSA-Decipher

In a typical PELSA experiment **(Figure 1)**, cell lysates extracted under native conditions are incubated with a ligand or vehicle, respectively. The two sample groups are then subjected to trypsinization with a high E/S ratio for a short time followed by removing large, partially digested protein fragments with an ultrafiltration procedure. The collected peptides are then analyzed by liquid chromatography-tandem mass spectrometry (LC-MS/MS) in data-independent acquisition (DIA) mode. The peptides are quantitatively compared between the two groups, and the peptide with the lowest p value among all quantified peptides of a protein is selected to represent its corresponding protein for target protein identification. Mapping the quantified peptides to protein sequences can generate local stability profiles, which reveals the protein regions responsive to the ligand binding. The dose-dependent local stability changes can also be assessed when PELSA experiments are performed using multiple ligand doses. Since the local stability changes of the target protein are dependent on the ligand binding occupancy, the dose that produces the half-maximal changes reflect the local binding affinity of the ligand for the corresponding protein segment. The dose-response local stability changes were termed as local affinity profiles. Therefore, from the PELSA experiments, we can obtain three types of information, i.e., the ligand binding proteins that are represented by the peptide with the lowest p value among all quantified peptides of a protein, ligand binding regions that are revealed by the local stability profiles, and binding affinity that are determined by local affinity profiles. Previously, all these parameters were determined by using R Scripts and required professional experience, which is not user friendly.

In this study, we present a specially designed software tool, PELSA-Decipher, to process and visualize the quantitative dataset generated by the PELSA experiment **(Figure 1B)**. PELSA-Decipher has three modules, i.e. differential analysis (DA), protein local stability analysis (ProLSA), and concentration-dependent analysis (CDA), to extract the information on ligand binding, i.e. binding proteins, the binding regions and the local affinity. The DA module utilizes the peptide-level quantification data for preprocessing and empirical differential analysis, which is performed using an empirical Bayes t-test. The differential results at both the peptide and protein levels are produced, and volcano profiles are generated to identify target proteins. Next, the peptide-level differential results from the DA module, along with protein domain information, are imported into the ProLSA module for further analysis. The generation of protein local stability profiles allows for the analysis of binding regions of ligand in proteins. The peptide quantification data from multi-concentration experiments can be analyzed by the CDA module. By fitting concentration-dependent curves at the peptide level and further refining the results, concentration-dependent curves and EC50 values are obtained. This enables the analysis of the binding affinity between each region of the protein and the ligand, as well as the overall protein-ligand affinity, which can be used for further validating the interaction between the protein and its ligand.

### PELSA-Decipher enables statistical analysis and sensitive identification of target proteins

The DA module in PELSA-Decipher utilizes a mathematical statistical model to identify differential proteins between experimental and control groups by applying predefined thresholds. By integrating peptide-level quantitative data, this module enables differential analysis at both the peptide and protein levels **(Figure 2A)**. The quantification data can be either from Spectronaut or DIA-NN. Take the PELSA results of staurosporine as an example, in which the quantitative data were obtained from Spectronaut.

**Figure 2.**
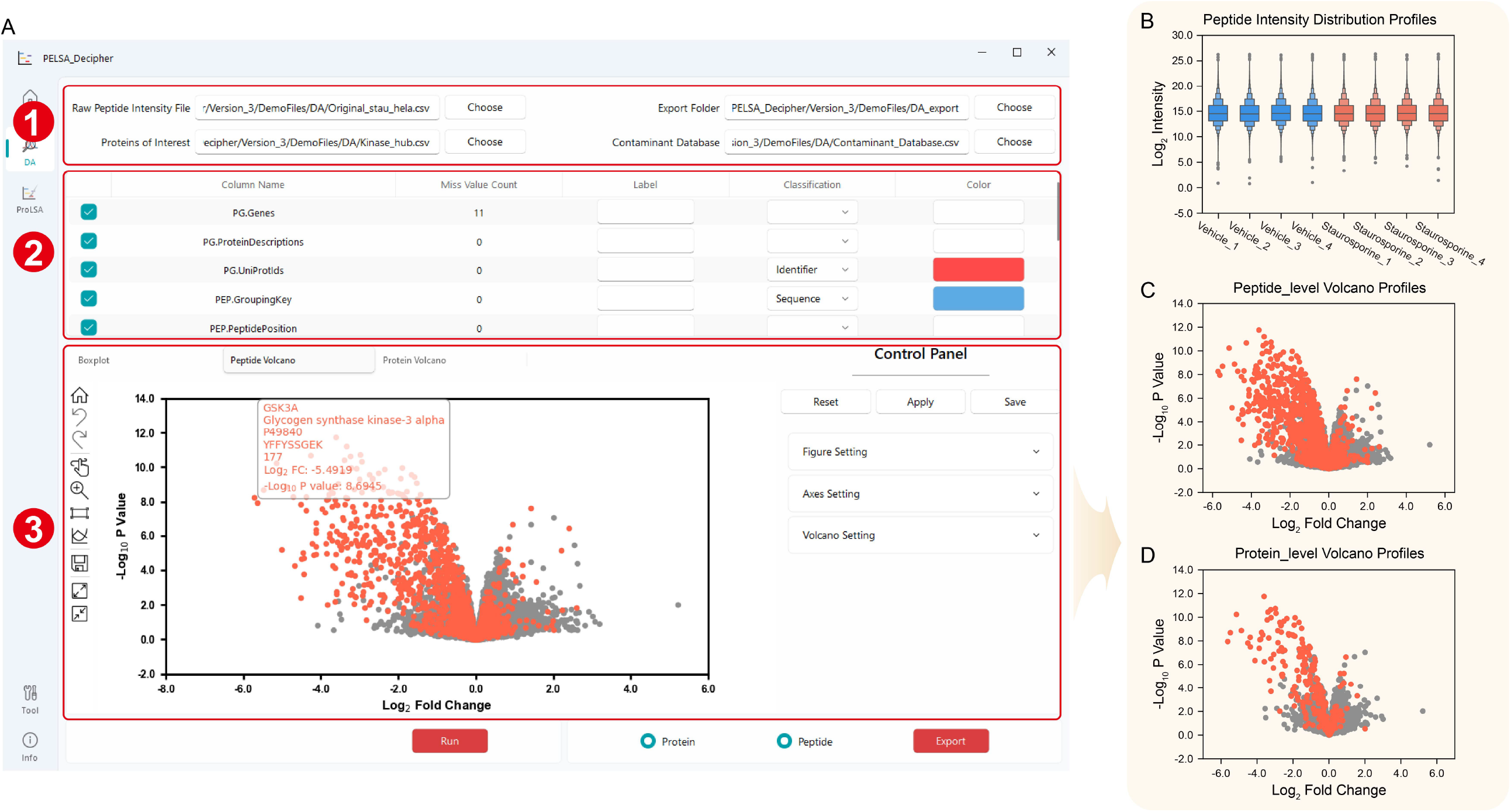
PELSA-Decipher enables statistical analysis and sensitive identification of target proteins. **A**. Interface of differential analysis. 1) Input and output file paths. 2) Parameter table for group classification. 3) Visualization of differential expression analysis results. **B**. The distribution of peptide intensity. Boxplot representation of log_2_ peptide fold changes. **C and D**. Peptide-level and Protein-level volcano plots, the orange point represent known target proteins of staurosporine. The data were obtained from HeLa cells treated with staurosporine using the PELSA method.

The experimental group was treated with staurosporine, while the control group was not treated, with four experimental replicates in each group. The acquired quantitative data exhibited high technical reproducibility, confirming the assumption that ligand-induced changes in proteolysis only affect a small subset of proteins in the proteome **(Figure 2B)**. Therefore, this procedure can be used to roughly evaluate the reproducibility of different replicates of PELSA experiment. Visualizing the distribution at both the peptide and protein levels using volcano profiles (with known target proteins highlighted in orange) revealed a clear stabilization trend in the target proteins and some of their peptides, consistent with the mechanistic principles of the PELSA approach **(Figure 2C, D)**.

To identify potential target proteins from quantitative dataset, some thresholds must be applied to filter candidate proteins. According to the principles of the PELSA method, proteins in the upper-left quadrant of the volcano profile are stabilized upon ligand binding, making them more likely to be potential target proteins. In contrast, proteins in the upper-right quadrant are destabilized by ligand binding, suggesting they might be indirect binding proteins that might be dissociated from targeting proteins when bounded by a ligand. By setting thresholds on both the -log_10_ p-value and log_2_ fold change, it is possible to effectively filter out potential target proteins. Given the readout amplification property of the PELSA method, target proteins typically exhibit significant fold changes, enabling the use of high threshold values (typically -log_10_ p-value ≥ 2, |log_2_ fold change| ≥ 1.5) to filter out the candidate binding proteins.

### PELSA-Decipher assists target protein filtering and binding region localization

The ProLSA module in PELSA-Decipher is designed to refine peptide quantitative data, which can be used to improve the accuracy of ligand-targeted protein filtering and enable the localization of binding regions. When ligands bind to proteins, the ligand bound regions of protein is more resistant to proteolysis, consequently, the intensity of target proteins differs markedly from that of non-target proteins, particularly in the vicinity of ligand-binding sites on the target proteins. Based on this phenomenon, constructing local stability profiles can facilitate the reliable determination of ligand-binding regions on target proteins and the candidate protein would be considered to be more reliable if obvious local stability was generated.

By integrating the peptide quantitative data with corresponding protein domain information, the amino acid sequence of the proteins is plotted along the x-axis, log_2_ fold change along the y-axis, and -log_10_ p-value is represented as a color gradient. Additionally, the structural domain intervals of the proteins are mapped to the x-axis. For ligand-target proteins, peptides near the binding sites typically exhibit log_2_ fold changes smaller than 0, indicating a tendency towards stabilization after ligand binding, while peptides located far from the binding sites generally show no significant changes (**Figure 3A**). Typically, the PELSA experiment can identify thousands of proteins, while PELSA Decipher can generate corresponding local stability maps for all of them. The obtain of this information could significantly enhances the efficiency and accuracy in determination of the reliable target proteins.

**Figure 3.**
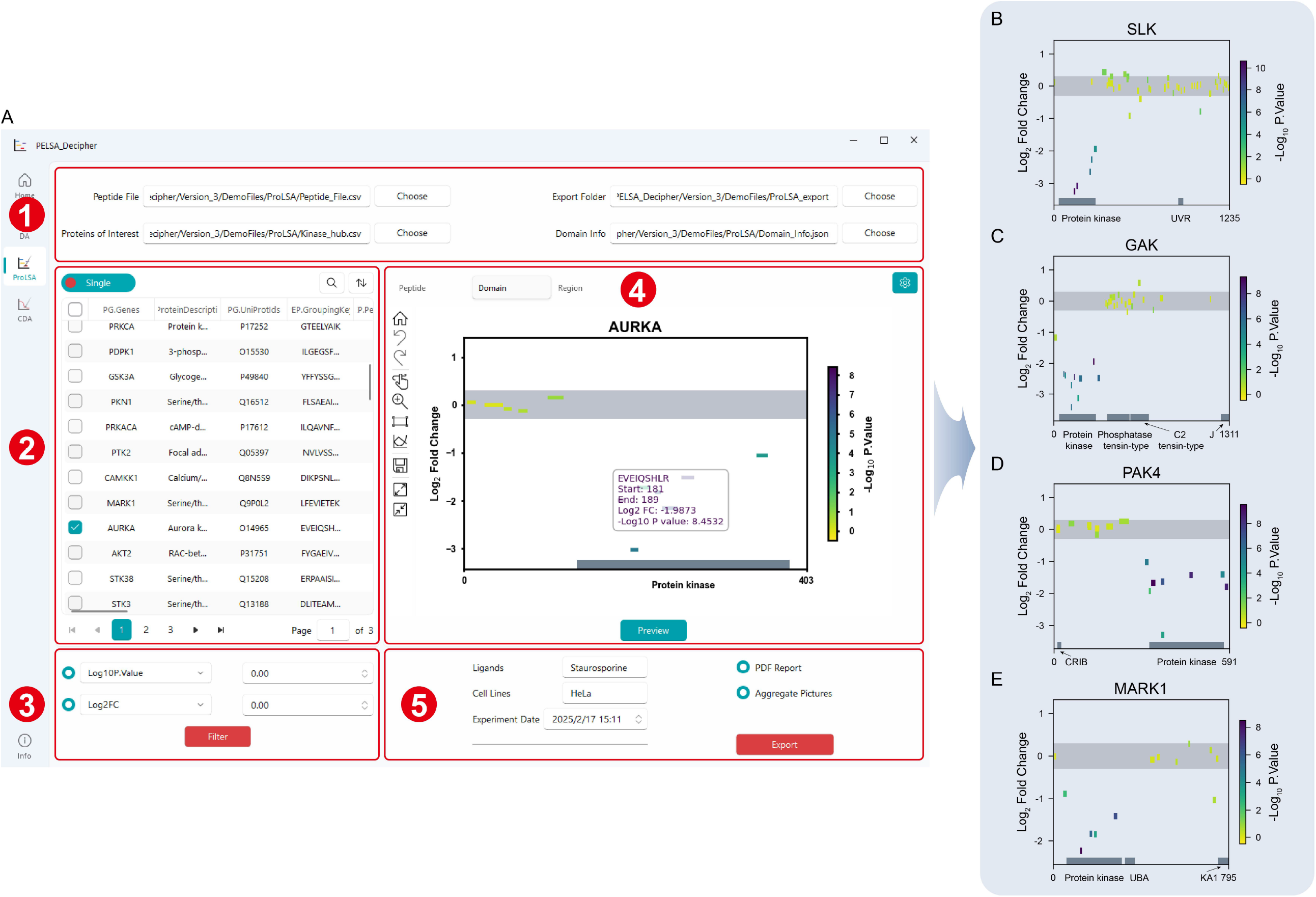
PELSA-Decipher enables the generation of local stability profiles for every target protein. **A**. Interface of protein local stability analysis. 1) Input and output file paths. 2) Table for selecting candidate proteins to preview local stability profiles. 3) Filter parameters to select proteins based on -log_10_ p-value and |log_2_ fold change|. 4) Display of protein local stability profiles. 5) Parameters for reporting output, including experiment information, date, PDF and image generation. **B, C, D and E**. Protein local stability profiles of staurosporine target proteins, SLK, GAK, PAK4 and MARK1. The x-axis represents the peptide sequence, the y-axis indicates the log_2_ peptide fold change, and the color bar represents the -log_10_ peptide p-value. Peptides in the Protein kinase domain exhibit significant stability trends, whereas peptides outside this domain show no stability differences. The data were derived from the PELSA analysis of staurosporine targets using lysates of HeLa cells.

Still use the PELSA results of staurosporine as an example. The known target proteins exhibited significant local stability in the protein-ligand binding regions, as shown by the protein local stability profiles **(Figure 3B, C, D, E)**. The four target proteins—SLK, GAK, PAK4 and MARK1—demonstrated consistent local stability, corresponding to the Protein kinase domain, as indicated by the domain information annotated in UniProt database. Detailly, the local stability profiles of these four proteins present a significant increase in peptide intensity (log_2_ fold change < 0, -log_10_ p-value > 2) in region corresponding to the kinase domain, while peptides outside the kinase domain, show no significant changes in intensity. For non-kinase proteins, such as CHD1L and RRBP1, all identified peptides show no significant changes in any sequence region, with most peptides falling within the range of |log_2_ fold change| < 0.3 **(Figure S1A, B)**. Above results suggest that the local stability profiles can assist in target protein filtering and binding region/site localization.

### PELSA-Decipher enables the determination of peptide-level binding affinity and protein-level binding affinity

The CDA module in PELSA-Decipher models the concentration dependence of peptides and proteins, enabling the calculation of both local and global affinities of ligands for their target proteins. By analyzing the magnitude of these affinities and the variations in local binding affinities, the CDA module facilitates the further assessment of target proteins **(Figure 4A)**.

**Figure 4.**
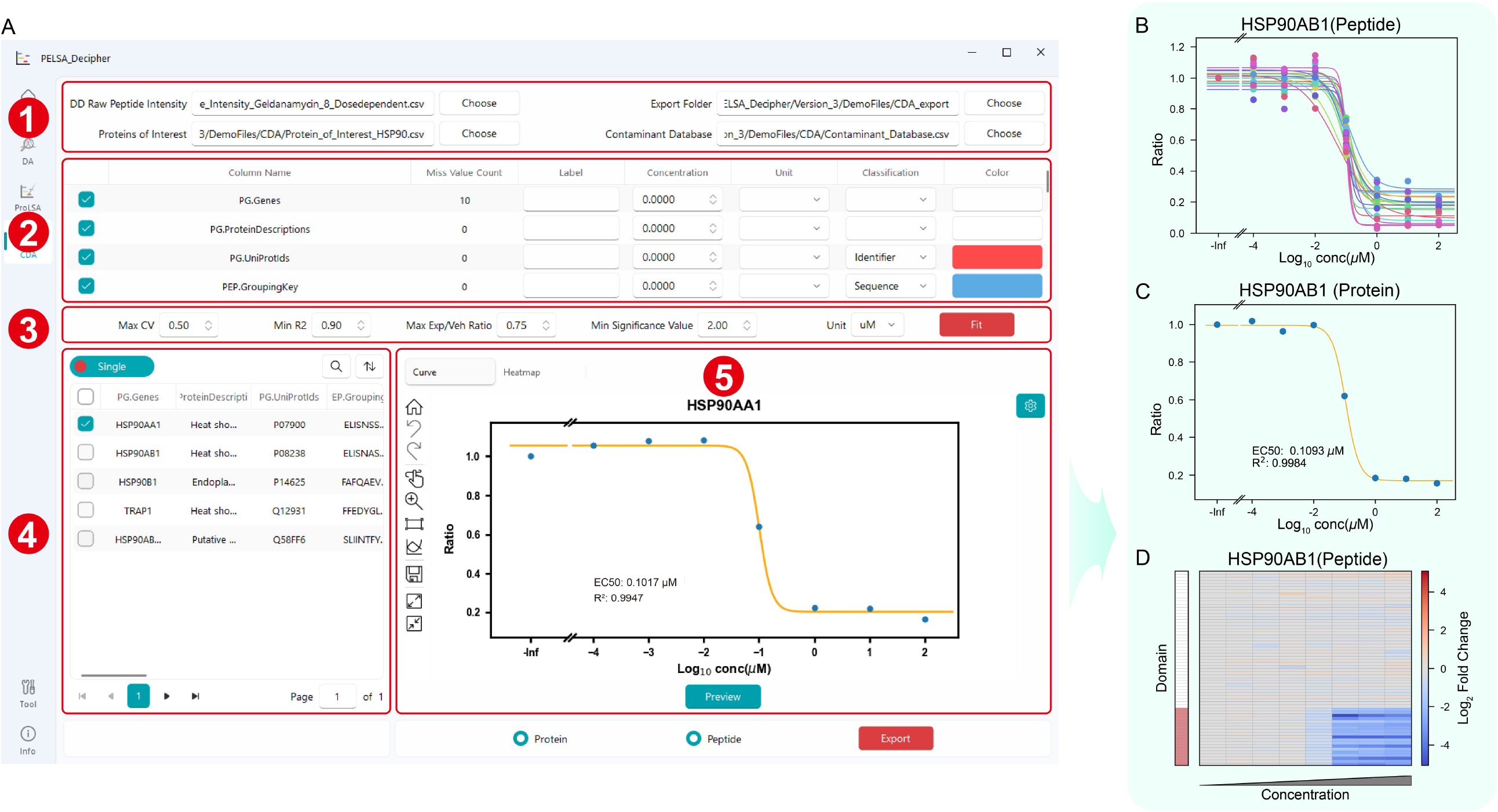
PELSA-Decipher enables the determination of peptide-level binding affinity and protein-level binding affinity. **A**. Interface of concentration-dependent analysis. 1) Input and output file paths. 2) Parameter table for setting group labels, concentrations, and classifications. 3) Parameters for filtering peptide data (max CV = 0.5) and selecting peptide fit data to calculate protein concentration dependence results (min R^2^ = 0.90, max Exp/Veh Ratio = 0.75, min significance value (−log_10_ p value) = 2.00). 4) Table for selecting candidate proteins to preview concentration dependence. 5) Display of protein and peptide concentration dependence curves and heatmap profiles. **B and C**. Concentration-dependent curves of HSP90AB1 peptides and protein. The EC50 value is 0.1093 μM, and the R^2^ value is 0.9984. **D**. Local affinity heatmap profile of HSP90AB1 with geldanamycin. Heatmap representation of log_2_ peptide fold change at various geldanamycin concentrations (0, 10−4, 10−5, 10−6, 10−7, 10−8, 10−9, and 10−^1^0 M). The data were obtained using the PELSA method from lysates of HeLa cells treated with multiple concentrations of geldanamycin.

Geldanamycin is a specific inhibitor of Hsp90 that binds to its N-terminal ADP/ATP binding site. In the geldanamycin experiment, eight concentration points—Vehicle, 10^−4^, 10^−5^, 10^−6^, 10^−7^, 10^−8^, 10^−9^ and 10^−10^ (M)—were used to measure the ligand-protein binding affinities. Taking HSP90AB1 as an example, at the peptide level, the identified peptides exhibited highly consistent binding affinities **(Figure 4B)**. At the protein level, a marked concentration-dependent effect was observed, with an EC50 value of 0.1093 μM **(Figure 4C)**. Heatmap visualization, combined with the mapping of domain information onto peptides, revealed that the stabilized peptides upon geldanamycin binding were predominantly located within the protein’s domain region (highlighted in red). These regions exhibited a significant concentration-dependent stabilization effect, providing further support into the confidence evaluation of ligand-protein interactions **(Figure 4E)**.

### PELSA-Decipher enables efficient and user-friendly data analysis

PELSA-Decipher, a professional and user-friendly software, significantly lowers the barrier to the data analysis of PELSA experiment, making the data processing more streamlined and efficient. Compared to original manual PELSA analysis[20], PELSA-Decipher achieves slightly better results in differential analysis and concentration-dependent analysis while markedly improving the processing efficiency. When analyzing staurosporine data, PELSA-Decipher (based on limma) achieves better result (108/135 versus 100/125) than the original processing (based on t-test) (**Figure S2A**). Additionally, in the fitting of 66,412 peptide concentration-response curves, PELSA-Decipher achieves a fivefold increase in processing speed compared to original processing based on R (**Figure S2B**). By leveraging protein local stability maps for target protein selection, users can generate and export results in batch with a single click, eliminating the tedious and repetitive steps of plotting and annotation.

## Discussion

Currently, numerous mass spectrometry-based screening techniques have been developed to identify ligand binding targets by detecting changes in protein’s property before and after ligand binding, as reflected in alterations in thermal stability, oxidative stability, and other related properties. Compared with other label-free methods, the PELSA approach demonstrates superior efficiency, sensitivity and precision, allowing for the accurate identification of ligand target proteins as well as the determination of their interaction regions. Given the substantial volume of dataset generated by the PELSA experiment and the complexity of its data analysis, we developed PELSA-Decipher. This tool can process large-scale PELSA data efficiently, providing multi-dimensional result visualization that can simplify target protein filtering and streamline the overall data analysis workflow. Additionally, we constructed protein local stability profiles for the PELSA method by combining experimental data with domain information to offer a more intuitive approach for target protein identification. The heatmap is used to display the ligand binding affinity calculated for each peptide, which is then mapped onto the corresponding protein domains. The combination of multiple level of PELSA information can assist the determination of the ligand binding proteins as well as the evaluation of the identification reliability.

Although PELSA Decipher is robust in data processing, it still has certain limitations. While it can also be used to handle data from other methods, its primary design is tailored for the PELSA approach, with a relatively fixed, standardized data processing workflow. As a result, adjusting specific processing steps can be somewhat challenging. Additionally, although PELSA Decipher is capable of data analysis and visualization, it currently does not provide a direct list of potential target proteins, functioning more as an auxiliary filtering tool.

In summary, PELSA Decipher is a data processing and result visualization software that enables efficient and accurate analysis of PELSA data, benefiting the identification of target proteins. Additionally, by adjusting the plotting parameters, it can generate the required visual outputs, facilitating further downstream analyses.

## Acknowledgments

This work was supported, in part, by funds from the National Key Research and Development Program of China (2021YFA1302600, 2024YFA1306000), the National Natural Science Foundation of China (22437007, 22137002), Dalian Science and Technology Innovation Fund (2023JJ11CG006), and United Foundation for Dalian Institute of Chemical Physics Chinese Academy of Sciences and the Second Hospital of Dalian Medical University (DMU-2&DICP UN202401).

## Notes

https://github.com/DICP-1809/PELSA-Decipher

### Competing Interest Statement

The authors have declared no competing interest.

