## Supplementary Figure Legends for "PELSA-Decipher: a software tool for the processing and interpretation of ligand-protein interaction dataset acquired by PELSA"

**Supplementary Figure 1: The local stability profiles of CHD1L and RRBP1.**

**A.** CHD1L. **B.** RRBP1.

**Supplementary Figure 2: PELSA Decipher has improved performance compared to traditional methods.**

**A.** PELSA Decipher employs the limma package in R for differential analysis, which, compared to the traditional t-test, is capable of identifying a greater number of target proteins at the same false positive rate. Specifically, PELSA Decipher identifies 108 kinases at an 80% false positive rate, whereas the traditional t-test yields only 100 kinases. This data is derived from the HeLa cell dataset within the PELSA method. **B.** PELSA Decipher, in conjunction with R, fits a total of 66,412 peptides. The time taken by PELSA Decipher to fit these peptides is 476 seconds. The complete process, including peptide fitting, data filtering, and final protein fitting, totals 494 seconds. In contrast, using R alone requires 2822 seconds and 2867 seconds, respectively, resulting in a fivefold increase in efficiency. This data is sourced from the Jurkat cell dataset within the PELSA method.
