## Supplementary figures and images for "PELSA-Decipher: a software tool for the processing and interpretation of ligand-protein interaction dataset acquired by PELSA"

### Figure S1

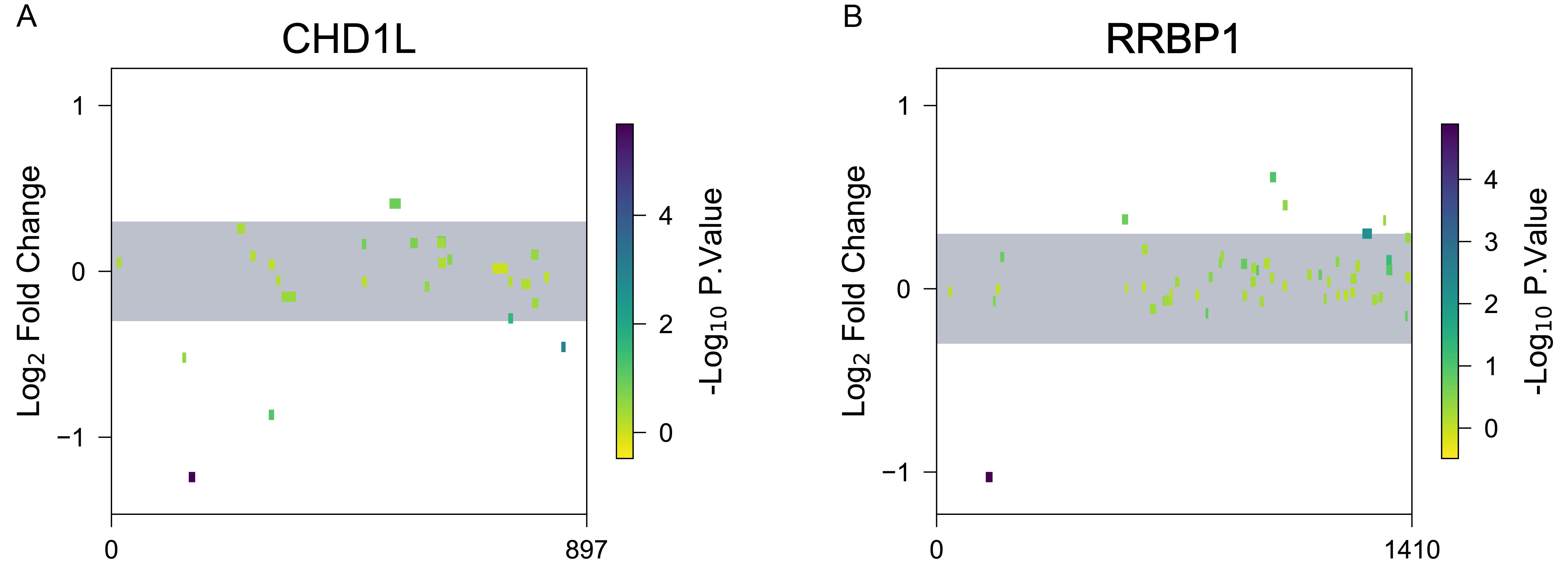

### Figure S2

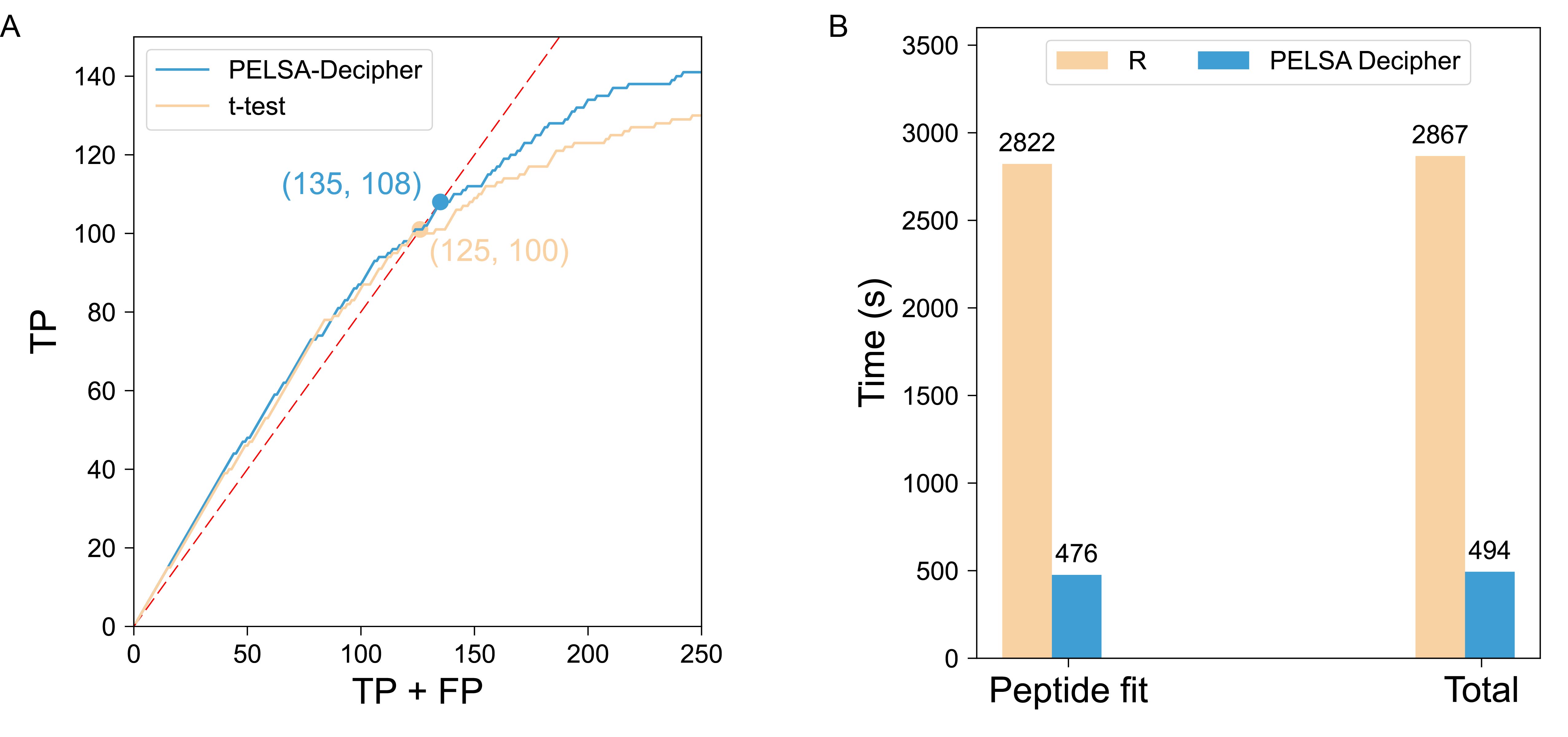
